# Tendon slack length is the primary determinant of plantarflexor muscle-tendon function in computational simulations of gait

**DOI:** 10.1101/328435

**Authors:** Josh R. Baxter, Michael W. Hast

## Abstract

***Background***: Locomotion is partly dictated by plantarflexor function and structure. Computational simulations are powerful tools capable of testing the isolated effects of muscle-tendon structure on gait function. ***Research Question***: The purpose of this study was to characterize the sensitivity of plantarflexor muscle function based on muscle-tendon unit (MTU) parameters. We hypothesized that plantarflexor metabolics and shortening dynamics would be sensitive to MTU parameters. ***Methods***: Stance phase of gait was simulated using a musculoskeletal model and computed muscle control algorithm. Optimal muscle fiber length, tendon slack length, and tendon stiffness parameters were systematically changed to test the effects on plantarflexor metabolics and shortening dynamics. ***Results and Significance***: Plantarflexor metabolic demands were 8 and 28 times more sensitive to muscle fiber and tendon slack lengths, respectively, compared to the effect of tendon stiffness. Shortened tendon slack lengths induced a large passive plantarflexion moment during early stance, which required non-physiologic dorsiflexor contractions. Conversely, longer muscle fiber and tendon slack lengths increased the shortening demands of the plantarflexors to account for the added length of the MTU. These findings highlight the importance of carefully selecting MTU parameters when modeling gait with musculoskeletal models, especially in pathologic or high-performance athlete populations.

## Introduction

Locomotor performance is explained, in part, by plantarflexor structure and function in both athletic and patient populations [1–4]. Longer muscle fibers allow sprinters to generate more joint power [5], which, in part, explains performance differences between good and great sprinters [6]. Tendon stiffness, a function of both its slack length and material properties, dictates the shortening demands of the plantarflexor muscles and impacts movement efficiency [7–9]. While these muscle-tendon parameters seem to explain functional differences in patient and athletic cohorts, the isolated effects of these structural measurements are difficult to elucidate due to variability inherent to *in vivo* research models.

Musculoskeletal models are powerful tools used with clinical gait analysis to test specific questions regarding muscle and tendon structure and function in isolated computational experiments. In these models, joints are actuated by muscle-tendon units (MTU) that have been developed using cadaveric measurements of muscle and tendon structure [10]. Patient-specific models are often scaled from these generic models but do not include subject-specific MTU parameters [11]. While these MTU parameters may be appropriate for some cohorts, special populations often have different muscle tendon structure [3,5]. Simulated gait mechanics are sensitive to MTU parameters [7,12–14], but the sensitivity of muscle shortening dynamics and metabolic requirements on these parameters are not fully understood.

The purpose of this study was to perform a sensitivity analysis on plantarflexor MTU parameters during simulated gait. Specifically, we utilized a musculoskeletal model to test the effects of plantarflexor fiber length, Achilles tendon slack length, and Achilles tendon stiffness on the energetics and muscle shortening dynamics of the stance phase of walking. These three parameters were selected because they have been documented to change in response to both training [15,16] and injury [17–19]. We hypothesized that (1) elongated tendons would lead to compromised function [3], (2) fiber length would be negatively correlated with energetic efficiency [7,20], and (3) tendon stiffness would be the strongest predictor of energetic efficiency [20].

## Materials and Methods

Human walking was simulated using open-source musculoskeletal modeling software (OpenSim, v3.3) and publically available gait data [11]. To reduce the complexity of human gait, we decided to utilize a computational model that moved only in the sagittal plane with simplified joints and a reduced set of muscles (Figure 1). The sagittal plane model consisted of 8 segments that had 10 degrees-of-freedom, defined by 7 sagittal-plane joints and actuated by 18 MTUs (gait10dof18musc, Opensim) and 3 pelvis degrees of freedom. The ankle was modeled as a pin-joint that was flexed by a single dorsiflexor muscle, the tibialis anterior, and extended by two plantarflexor muscles, the soleus and gastrocnemius. Notably, the soleus and tibialis anterior muscles are uniarticular, meaning that an ankle motion is achieved when the muscle is flexed. Conversely, the gastrocnemius is biarticular, so coupled plantarflexion of the ankle and flexion of the knee flexion when this muscle is flexed. Muscles were modelled as Hill-type muscle bundles (Millard type [21]) that included a contractile muscle element in series with an elastic tendon-like element. The structural parameters of the MTU included optimal fiber length, tendon slack length, and tendon stiffness defined as the amount of tendon strain experienced under isometric maximal muscle force (**Table 1**).

**Figure 1.**
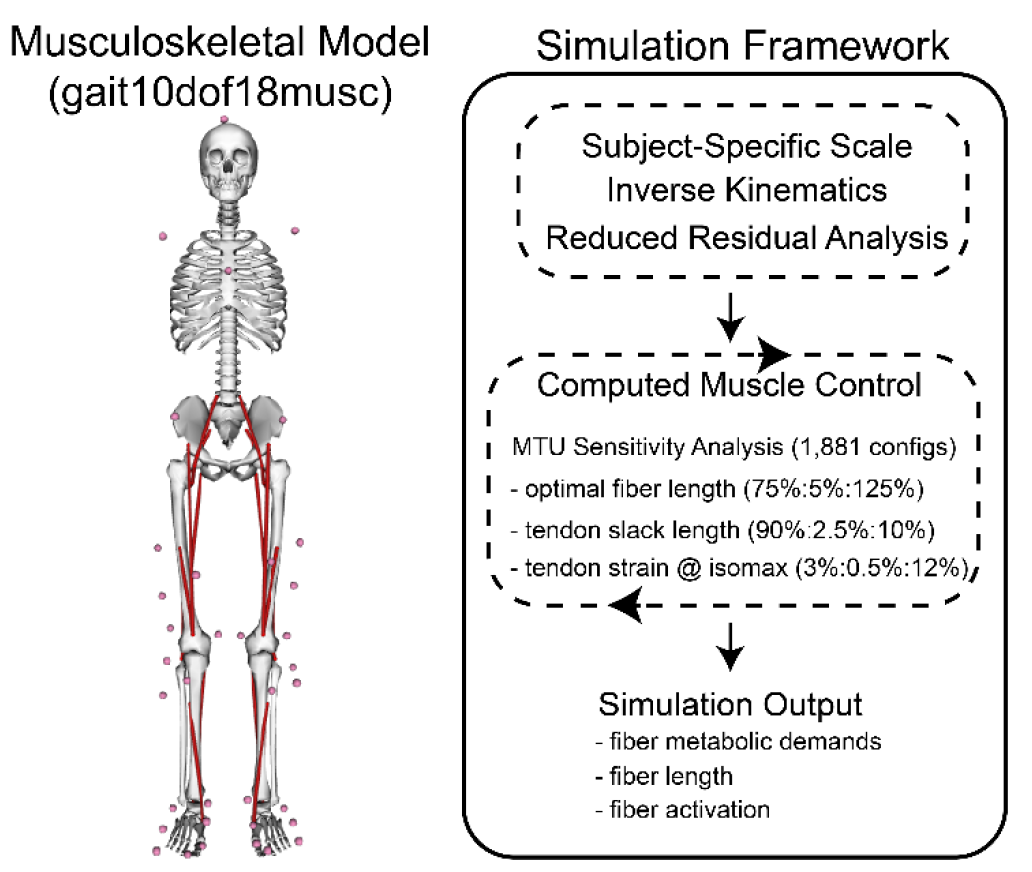
Stance Phase of gait was simulated using a musculoskeletal model constrained to move in the sagittal plane (*left*). A sensitivity analysis was performed on three muscle-tendon unit(*MTU*) parameters: optimal fiber length, tendon slack length, and tendon strain at isometric fiber force (*right*). 1,881 simulation outputs were compared using a regression model.

**Table 1.** Default muscle tendon unit (MTU) parameters.

|  | Gastroc. | Soleus |
| --- | --- | --- |
| Max Isometric Force | 2500 N | 5137 N |
| Optimal Fiber Length | 9.0 cm | 5.0 cm |
| Tendon Slack Length | 36.1 cm | 25.1 cm |
| Tendon Strain @ Isomax | 4.9% | 4.9% |

Stance phase during gait was simulated using a previously established framework [11] (**Figure 1**). Skin-mounted marker trajectories and ground reaction forces of a healthy-young male (1.80 m, 75 kg) who walked at a comfortable speed (1.2 m/s) were utilized in this study, which was provided with the musculoskeletal modeling distribution (Gait10dof18musc\ExperimentalData). First, a generic model was scaled to match the anatomic proportions and body mass of the research subject. Second, a time series of joint angles was calculated to best fit the experimentally acquired motion data. This data satisfied the joint definitions of the sagittal plane model and was calculated using the on-board Inverse Kinematics algorithm. Third, the Residual Reduction algorithm was performed to adjust the torso model mass and joint coordinate trajectories to ensure dynamic consistency of the simulation. Finally, the Computed Muscle Control (CMC) algorithm [22] calculated muscle excitations that generated model motion that matched the kinematics and externally applied forces while minimizing the sum of squared muscle activations.

A sensitivity analysis was performed to characterize the effects of optimal muscle-fiber lengths, Achilles tendon slack lengths, and Achilles tendon stiffness variations (**Figure 1**) on muscle metabolic demands and shortening dynamics. Fiber lengths of the ankle plantarflexors (soleus and gastrocnemii) were simultaneously scaled in 5% increments from 75% to 125% of the standard model lengths. Achilles tendon slack lengths were scaled values of the model default values, from 90% to 110% of original length in 2.5% increments. Achilles tendon stiffness values were modeled as the tendon strain at the maximum isometric force of each plantarflexor muscle, which ranged from 3 to 12% in 0.5% strain increments. The dorsiflexor muscle parameters were not manipulated. In total, this parameterization study tested 1,881 combinations of muscle fiber lengths, Achilles tendon slack lengths, and Achilles tendon stiffness values were computed.

Metabolic demands and shortening dynamics of the ankle muscles were evaluated during each simulation to quantify the effects of varying muscle-fiber lengths, tendon slack lengths, and tendon stiffness. Metabolic energy requirements of the gastrocnemius, soleus, and tibialis anterior muscles were estimated using a metabolic calculator that predicted thermal and mechanical energy liberation of muscle contractions [23] that were normalized by body weight. Muscle activations, shortening velocities, and active fiber powers as well as tendon powers were directly acquired from the musculoskeletal modeling software.

The effects of MTU parameters on muscle metabolics were quantified using a linear regression model (MATLAB, Mathworks, Natick, MA). Some simulations could not run to completion because of passive plantarflexion moments that were too large to be countered by active dorsiflexor contractions. Because of this, we also performed multivariate linear regression to explain the effects of MTU parameters on dorsiflexor (tibialis anterior) metabolics. The effect sizes of each MTU parameter was reported as the change in metabolic energy consumption per kilogram of bodyweight for a 1% change in MTU parameters compared to the model default values.

## Results

Fiber length and tendon slack lengths had effects of 8 and 28-fold greater on plantarflexor metabolic demands than tendon stiffness, respectively (**Table 2**). Despite its smaller effect size on metabolic demands, tendon stiffness demonstrated a parabolic effect on metabolics (**Figure 2**). Conversely, both muscle fiber and tendon slack length imposed additional metabolic demands with increased length. These effects were explained by increased muscle shortening demands that were required to take the slack out of the MTUs. Plantarflexor muscle activation patterns were most sensitive to changes in MTU parameters during late stance (**Figure 3A/C**), particularly fascicle and tendon slack lengths (**Figure 3 top and middle row**). Similarly, longer muscle fibers and tendon slack lengths required greater plantarflexor shortening in order to maintain the necessary tendon tension (**Figure 3B/D**).

**Figure 2.**
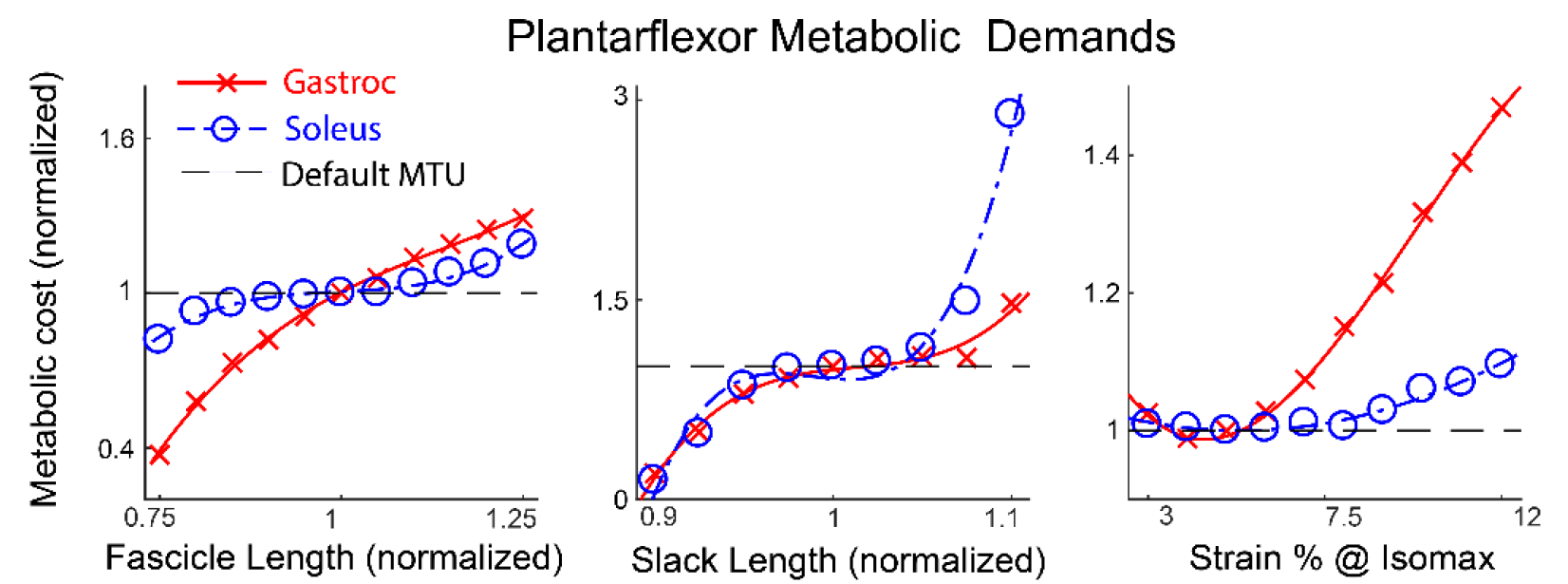
Small changes in muscle tendon unit(*MTU*)parameters (*X-axis*) strongly affected metabolic demands of both plantarflexor muscles (gastroc-blue crosses, soleus-orange circles). Metabolic demands (Y-axis) were normalized to the model default MTU parameters(dashed line).

**Figure 3.**
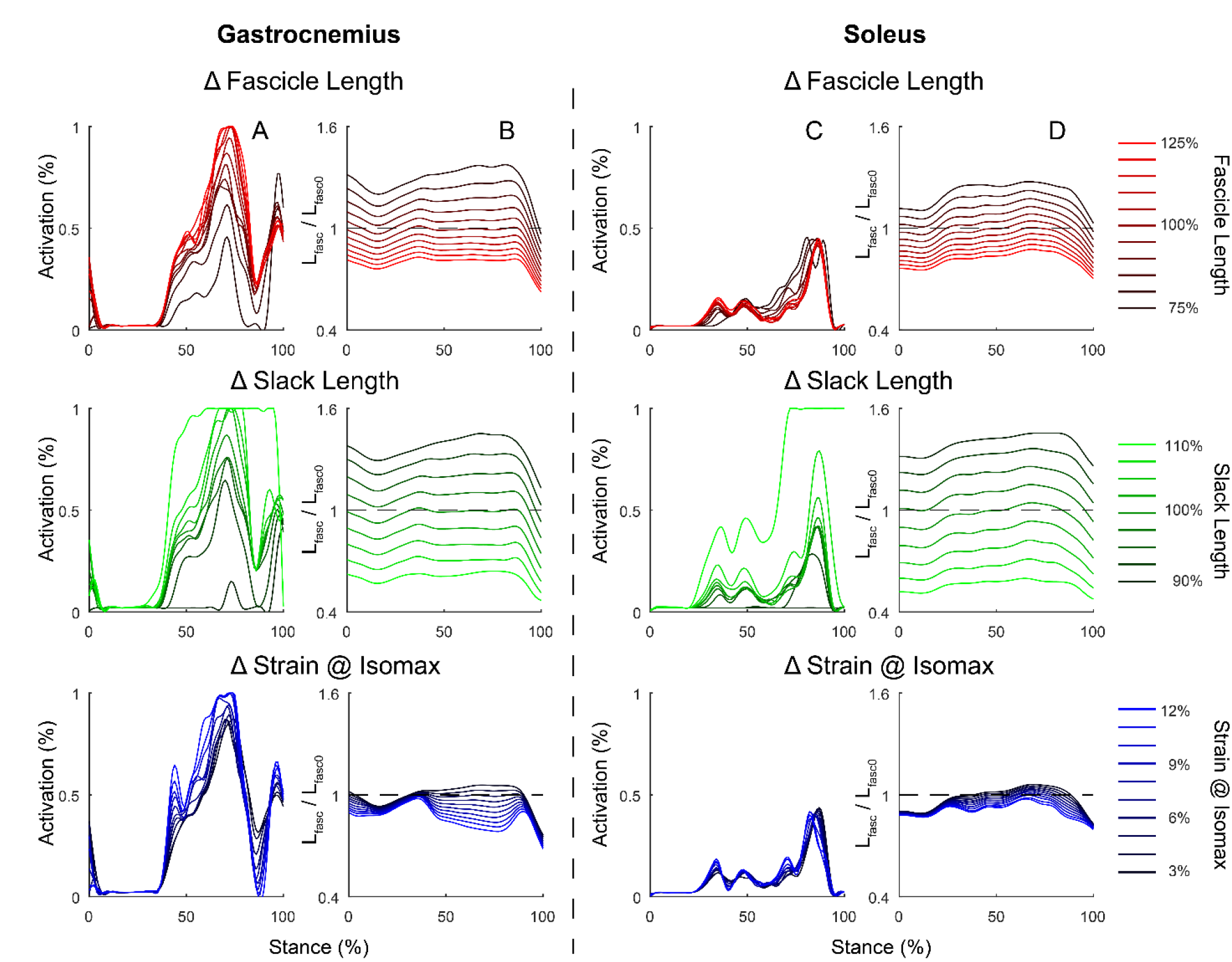
Plantarflexor activations (*A,C*)and fascicle lengths (*B,D*) during stance phase were more sensitive to changes in optimal fascicle length (*top row,red*) and tendon slack length(*middle row, green*) that tendon compliance (*bottom row,blue*). Fascicle length during stance phase was normalized by the optimal fibre length for that specific simulation(L_fasc_/L_fasc0_). For clarity, these plos demonstrate the isolated effects of muscle tendon unit(*MTU*) parameters on activation and fascicle length change by holding the other MTU parameters at their defaults (Table 1).

**Table 2.** Effect of 1% change of MTU parameter on PF and DF metabolics *P < 0.001* for all variables.

| | $\Delta$ PF metabolics | $\Delta$ DF metabolics |
| --- | --- | --- |
| Fascicle length | 2.01% | -0.53% |
| Tendon slack length | 6.69% | -2.99% |
| Tendon compliance | 0.23% | <0.01% |

*P < 0.001* for all variables.

Tendon slack length had a strong effect on dorsiflexor metabolic demands during stance phase of gait (**Table 2**). This demand was so great that a 10% decrease in the plantarflexor tendon slack lengths resulted in simulations that could not run to completion when fiber lengths were less than 90% of the default model values. While longer muscle fibers helped the simulations run to completion, the metabolic costs of the dorsiflexor muscle were 2–10 times greater than the default MTU parameters until muscle fibers were set to at least 15% longer than default, regardless of tendon stiffness.

Plantarflexor metabolic demands were minimized when the MTU was short (fiber length and slack lengths 80% and 92.5% of default, respectively) and the tendon was stiffest (3% strain at isometric max). This metabolic minimum was explained by a short and stiff MTU that passively stored and returned energy during midstance (**Figure 4C**), which reduced the amount of plantarflexor shortening and activation needed in late stance (**Figure 4A/B**). However, this came at the expense of increased dorsiflexor metabolic costs of 10-fold greater than the energetic needs of the default MTU parameters. Additionally, MTU power was 2–3 times greater (**Figure 4D**) in this ‘optimal’ MTU configuration, which demonstrates that this Because these large metabolic demands were non-physiologic, simulations that required more than 150% of the default dorsiflexor metabolic demands were excluded from all analysis. After excluding these simulations, plantarflexor metabolic demands were minimized when the muscle fibers were their shortest (75% of default), the tendon slack lengths were set to their default lengths, and the tendons were stiffest (isometric muscle force at 3% strain).

**Figure 4.**
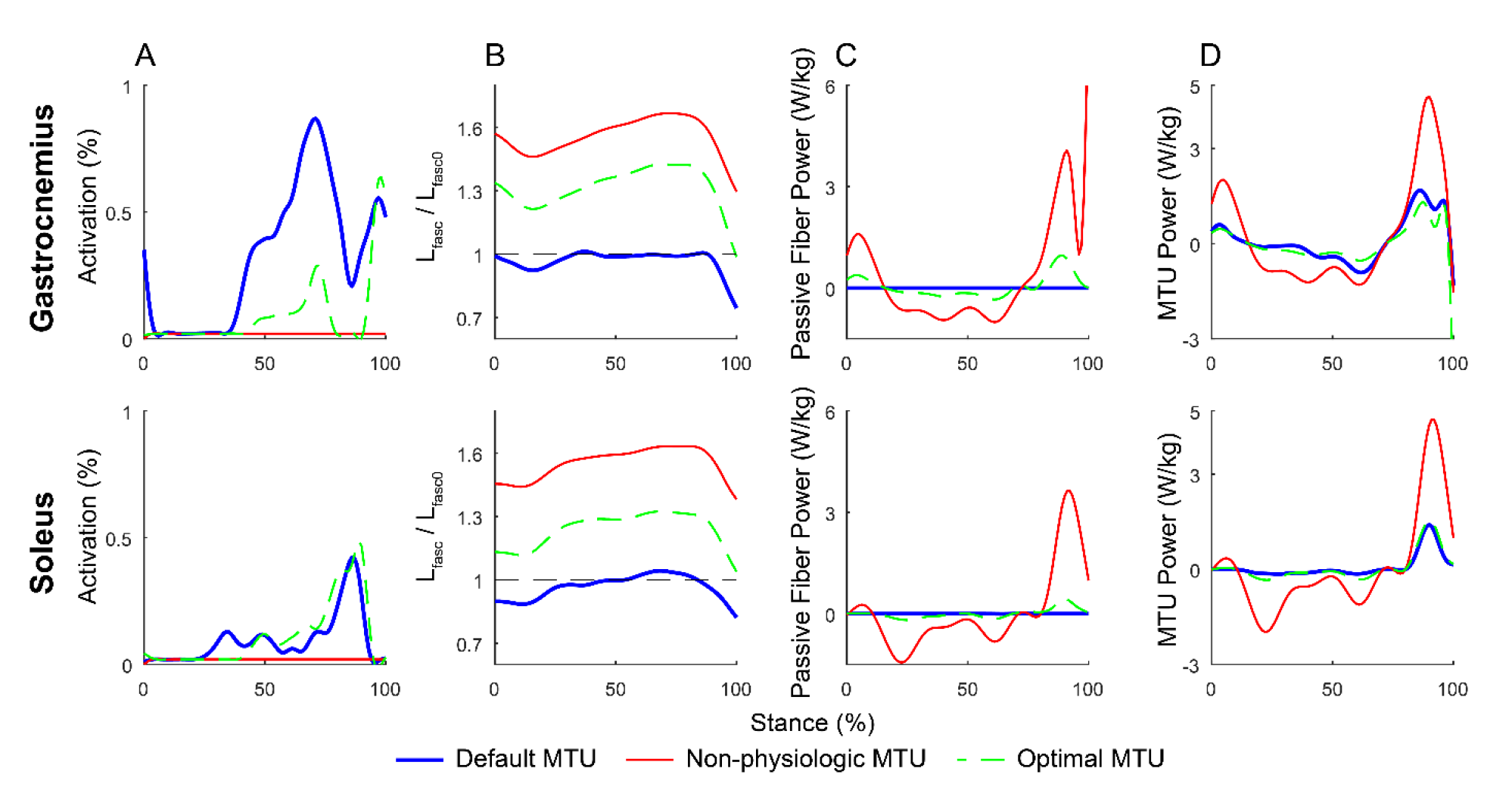
Muscle activation (*A*), normalized fiber length (*B*), passive fiber power (*C*), and MTU power (*D*) differed between optimized plantarflexor shortening dynamics and the defafult MTU parameters (*thick blue line*). Non-physiologic MTU parameters(*red line*) resulted in very short MTUs that required no activation to generate large amounts of passive fiber power. Optimized MTU parameters(*dashed green line*), within physiological conditions, reduced the gastrocnemius (*top*) activations but not the soleus(*bottom*) activation patterns.

**Figure 5.**
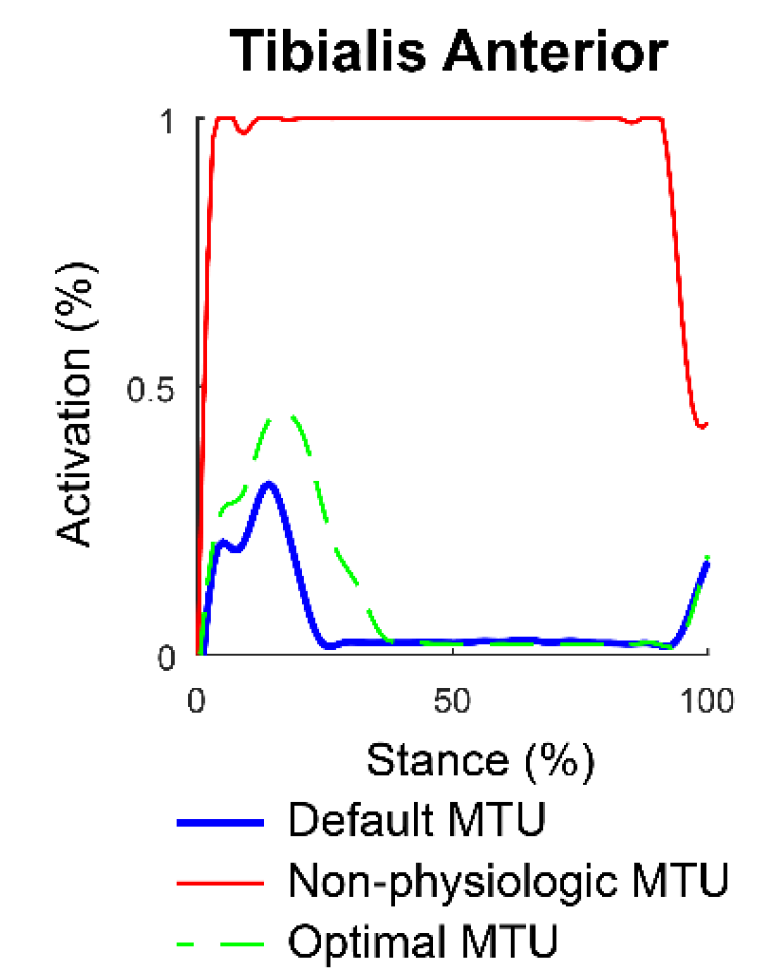
Tibialis anterior activations were maximal with non-physiologically short fiber and tendon slack lengths(*red*). Optimized MTU parameters(*dashed green line*),within physiologic conditions, required slightly greater dorsiflexor activation compared to the default MTU simulation(*thick blue line*).

## Discussion

Musculoskeletal simulations are powerful tools for studying clinically important questions. In the present study, we highlighted the sensitive relationship between MTU parameters and muscle shortening dynamics and metabolic demands. The resting length of the MTU, dictated by fiber and tendon slack lengths, had profound effects on plantarflexor shortening dynamics and metabolics (**Figure 2** **and 3, Table 2**). Surprisingly, tendon stiffness was far less influential, despite the substantial relationships between stiffness and metabolics that has been reported by others [7,8]. This discrepancy may be explained by experimental evidence suggesting that fiber lengths undergo remodeling in response to changes in tendon slack lengths in order to maintain tendon tension for some activity [24], joint posture [25,26], and tendon injury [27,28]. The results of the current study highlight the importance of MTU parameters on the metabolic cost of locomotion, which should be considered carefully when simulating human motion.

Simulation results reported in this study compare favorably with previous computational investigations on the effects of MTU parameters on locomotor function. Tendon slack length is the strongest predictor of gait mechanics, which agrees with several other sensitivity studies [12,13,29,30]. Tendon stiffness had a parabolic effect of plantarflexor metabolics, with optimal muscle function occurring at 4.5 – 5% strain at maximum isometric force, which matches the a previous report that leveraged the same inputs [8]. Plantarflexor metabolic demands were minimized when the muscle fibers were near their shortest lengths, which agrees with a prior report that found short muscle fibers in series with compliant tendons were optimal for walking [7]. However, we found that stiffer tendons were more energetically efficient, which may be explained by differences in muscle models [21] or control schemes [22]. Stiffer tendons reduced fiber shortening during the second half of stance (**Figure 3B/D**), suggesting that strain energy can be more efficiently stored with less active muscle shortening.

Shorter fascicles improve metabolic efficiency (**Figure 2**) and walking speed [20], which is in conflict with reports of longer muscle fibers being advantageous for faster motions like sprinting [5,31]. This difference between walking and running may not be that surprising given that walking is an energy efficient activity while sprinting is governed by how much work can be done to the ground. However, activities of daily living like stair climbing and rising on the toes are challenging in patients who have suffered Achilles tendon injuries, whom often develop elongated tendons [3]. Thus, there is a dichotomous need for shorter muscle fibers in order to walk efficiently and longer fibers to perform other tasks. Exercise prescriptions have been shown to improve muscle and tendon structure as a means towards improving function in adults [15,32–34].

Based on our findings, simulation results are very sensitive to MTU parameters (**Table 2**) and must be carefully considered when utilizing musculoskeletal models to study muscle and tendon function. One approach is to ‘tune’ the MTU based on experimental data collected *in vivo* or cadaveric donors [35]. In this scenario, each of the joints are set to a desired position, the muscles are excited with a nominal excitation, and the tendon slack length is subsequently changed to ensure the fiber length is set to its optimal length when the MTU is in equilibrium. Unfortunately, this approach relies on resting joint angles that may vary between patient cohorts and within individuals [25]. Alternatively, maximal muscle contractions can be simulated and the model can be ‘tuned’ to match experimental data [36]. This approach also has limitations, as it relies on the assumption that a voluntary maximal isometric contraction is equivalent to a true maximal contraction. A final option consists of tuning MTU parameters based on medical imaging [3] and *in vivo* methods [17,37] to provide subject-specific inputs. While this may seem ideal, this approach is time consuming and financially burdensome.

Several limitations should be considered when discussing the implications of this work. Model complexities were reduced to improve simulation speed, while reducing the uncertainty introduced by additional degrees of freedom and MTU actuators. Determining the appropriate control scheme for simulating human motion is challenging. We decided to utilize the Computed Muscle Control algorithm that minimized the sum of squared muscle activations [22,38]. This approach is reasonable for simulating the motions and interaction forces of the body during walking, and adequately characterizes the required muscle dynamics to generate a consistent walking pattern. However, this approach is likely not suitable for high-intensity activities, where minimization of muscle forces is not appropriate. The plantarflexor muscles were modeled as two separate muscles with independent tendons rather than a combined tendon that represented the Achilles tendon. Although the Achilles tendon is often recognized as a single connective tissue at the confluence of the plantarflexor muscles [CIT], recent evidence suggests that the discrete bundles that comprise the Achilles tendon function differently during walking [39]. Finally, we decided to utilize a well-documented and freely available data set so others can repeat these simulations; however, other activities could easily be implemented in future sensitivity analyses.

## Conclusions

We performed a parameterization study to better understand the effects of MTU parameters on plantarflexor shortening dynamics and metabolic and during gait. The sensitivity of plantarflexors MTU parameters were tested due to this muscle group’s effect on function in both athletic [6], pathologic [3], and aging populations [2,20]. Our findings demonstrate that simulation results are very sensitive to muscle fiber and tendon slack lengths, which should be carefully selected based on either experimental or optimization approaches. Ongoing work is focused on improving MTU tuning algorithms aimed at modeling more physiologic muscle-tendon function.

## Acknowledgments

The Authors have no acknowledgements

## Authors’ Contributions

JB and MH designed the experiment; JB performed the simulations and analyzed the data; JB and MH analyzed and interpreted the data; JB drafted the manuscript; JB and MH revised the intellectual content of the manuscript; JB and MH approved the final version of the manuscript; and JB and MH agreed to be accountable for all aspects of the study.

***Funding*** no funding has been provided for this research

